# Client recognition differences between PDI and ERp46 to guide oxidative folding

**DOI:** 10.1101/2024.03.04.583432

**Authors:** Tomohide Saio, Kotone Ishii, Motonori Matsusaki, Hiroyuki Kumeta, Shingo Kanemura, Masaki Okumura

**Affiliations:** Institute of Advanced Medical Sciences, Tokushima University, 3-18-15 Kuramoto-cho, Tokushima, 770-8503, Japan; Frontier Research Institute for Interdisciplinary Sciences, Tohoku University, 6-3 Aramakiaza Aoba, Aoba-ku, Sendai, Miyagi 980-8578; Global Station for Soft Matter, Global Institution for Collaborative Research and Education, Hokkaido University, Kita 21 Nishi 11, Kita, Sapporo, 0110021, Japan

## Abstract

Endoplasmic reticulum (ER)-resident protein disulfide isomerase (PDI) family members function not only as disulfide bond-catalysts but also as chaperones for the oxidative folding of client proteins. However, due to the scarcity of structural data, the client recognition mechanism remains poorly understood. We report the distinct recognition mechanisms for PDI/ERp46 proteins to recruit a reduced and denatured bovine pancreatic trypsin inhibitor (BPTI). NMR data demonstrated that PDI recognizes a broad region consisting of ∼30 amino acid residues in unfolded BPTI in order to function as a chaperone, while ERp46 interacts nonspecifically and weakly with unfolded BPTI to promiscuously and rapidly introduce disulfide bonds. These two distinct client recognition modes are likely important for the functional modulation of the disulfide-catalyst and chaperone against client proteins. Client recognition differences between PDI family proteins likely contribute to determining whether disulfide bonds are introduced into the client protein or aggregates are reduced to an unfolded state during the early folding step. Thus, the mammalian ER ensures concerted protein quality control through the differences in the client recognition modes of PDI family proteins, resulting in the production of large quantities of multiple-disulfide bonded proteins.

## Introduction

Protein folding is conformational convergent process to native conformation but intrinsically error-prone, and thus it must be guided along a funnel-shaped energy landscape involving multiple folding pathways to achieve a unique, stable, and native conformation^1^. Off-pathway folding intermediates expose unfavorable hydrophobic surfaces that can engage in aberrant intra- or inter-molecular interactions that enable aggregation^2^. To avoid the risk of proteotoxic aggregation, cells harbor complicated but sophisticated foldase/holdase networks that control protein folding to maintain protein homeostasis (proteostasis). Previous research into the proteostasis system of the cytosol has identified many chaperone proteins, such as heat shock proteins, which are induced by several signaling pathways, some of which are stress responsive, to ensure protein folding^3^. Despite the fact that chaperones represent a common recognition features of client sequences enriched in hydrophobic residues, different families of chaperones exhibit distinct activities against clients. Structural information on the complexes formed between chaperones, trigger factor (TF)/SecB, and clients has helped elucidate the different modes of client recognition; TF accommodates a stretch of up to 50 interacting residues to function as a foldase^4^, while SecB accommodates 250 interacting residues to function as a holdase^5^. Although structural information on chaperone-client complexes has been reported, the amount of information is limited and, importantly, no unifying mechanism on how chaperones recognize non-natively folded clients to guide folding has been reported.

Besides cellular protein folding by cytosolic chaperones, many nascent polypeptides are inserted into the endoplasmic reticulum (ER), where oxidative folding, coupled with disulfide bond formation occurs. More than 20 members of the protein disulfide isomerase (PDI) family are known to function in the ER as both chaperones and disulfide bond-catalysts^6–9^. Although PDI family members have different amino acid sequences, they all contain at least one thioredoxin (Trx)-like domain with or without a redox active CxxC motif, and the three-dimensional arrangements of these Trx domains are distinct from one another, resulting in different functionalities^7,9–16^. For instance, ERp46 engages in rapid but promiscuous disulfide bond formation while canonical PDI acts as a versatile catalyst for the introduction of native disulfide bonds into client proteins^16^. To obtain a deeper understanding of the mechanism underlying client recognition by PDI, structural determinations by us and others have shown that the oxidized form of PDI is in rapid equilibrium between open and closed U-shaped conformations, whereas the reduced form maintains a closed U-shaped conformation^17,18^, Regarding the client binding/release by ERp46, both oxidized and reduced ERp46 form an opened V-shaped conformation via two unusually long loops between Trx-like domains^19^ (Figure 1a) suggesting that ERp46 redox active Trx-like domains act independently and engage in rapid but promiscuous disulfide bond formation during early oxidative folding. Conversely, PDI catalyzes native disulfide bond formation via the cooperative action of two redox active sites on client folding intermediates bound to the central cleft of the U-shape conformer. Despite the accumulation of structural insights into PDI/ERp46-mediated protein folding, it is currently unclear how PDI and ERp46 recognize unfolded clients during the early stage of client folding.

**Figure 1.**
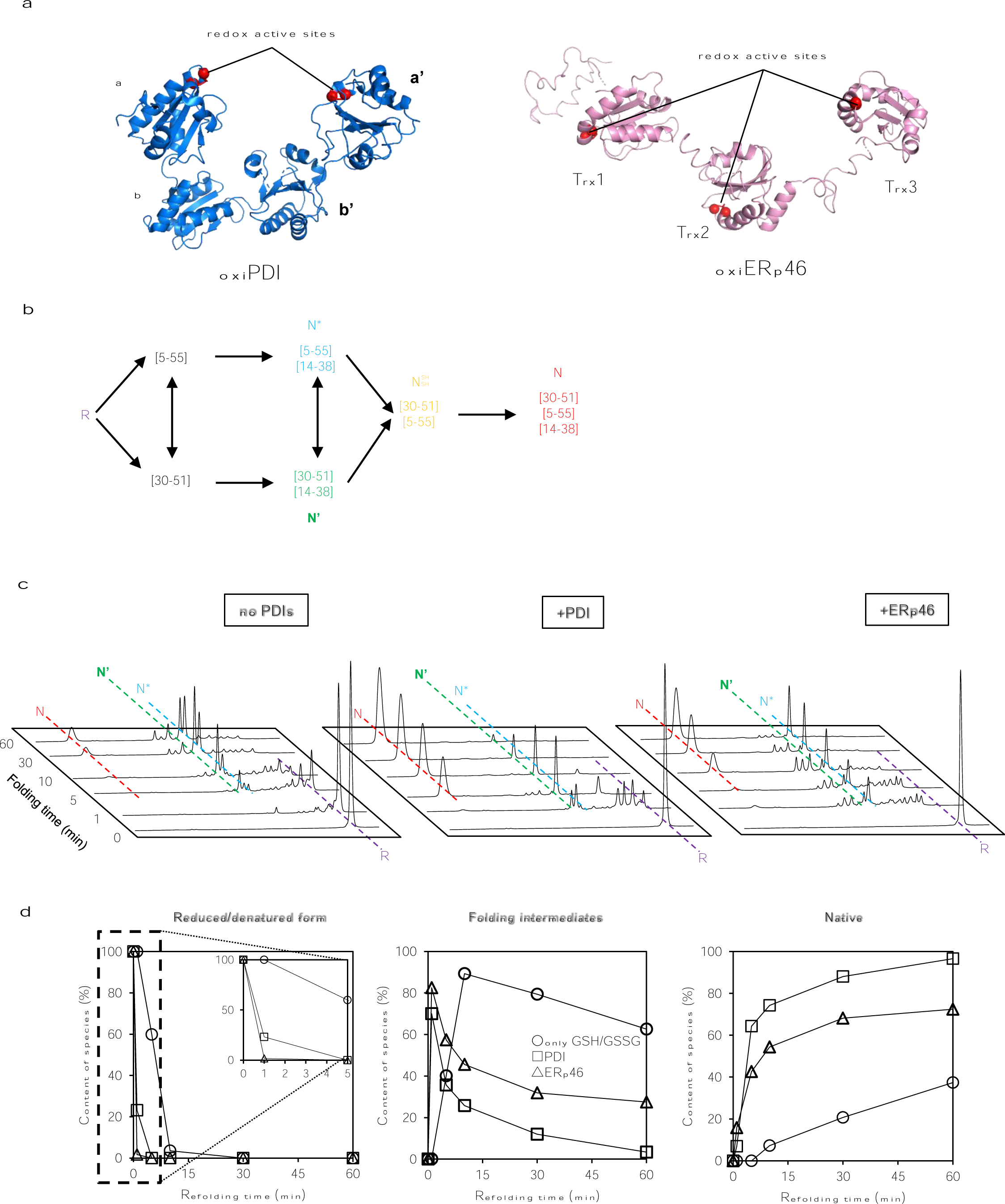
Oxidative folding of BPTI catalyzed by PDI and ERp46. (a) Overall structures of PDI (PDB: 4EKZ) and ERp46^19^. (b) Schematic representation of oxidative folding pathway of BPTI. R and N indicate reduced and native BPTI [Cys30-Cys50, Cys5-Cys55, and Cys14-Cys38], respectively. (c) HPLC profiles of the time course of oxidative folding by PDI and ERp46 in the presence of 2 mM GSH and 1 mM GSSG. N, N’, N*, NSH, and R indicate the disulfide bond patterns shown in Figure 1b. (d) Quantitative analyses of the percentages of reduced BPTI, folding intermediates, and native BPTI as a function of incubation time.

Here we employed a bovine pancreatic trypsin inhibitor (BPTI) mutant, which mimics a random coil in the initial stage of oxidative folding, and observed differences in the recognition of BPTI by PDI and ERp46 using NMR. Strikingly, PDI interacted with a broad region from residues 7 to 40 within the unfolded BPTI and is highly effective in inhibiting aggregation; however, the introduction of disulfide bonds by PDI is slower than that of ERp46. ERp46 interacts more weakly with unfolded BPTI than PDI without engaging in residue-specific client recognition for the rapid introduction of disulfide bonds into BPTI. It is thus conceivable that the client recognition differences between PDI family members in the mammalian ER ensures the concerted protein quality control needed for the production of large quantities of secretory and membrane proteins.

## Results and Discussion

### PDI/ERp46-catalyzed oxidative folding

BPTI consists of 58 amino acid residues and harbors three disulfide bonds (Cys5-Cys55, Cys14-Cys38, and Cys30-Cys51) in the native folded conformation. The BPTI oxidative folding pathway mediated by GSH/GSSG has been well characterized by Weissman and Kim ^20^(Figure 1b). When we previously evaluated the protein folding functions of PDI and ERp46 using BPTI as a model substrate^15^, we used 2 mM GSH and 0.2 mM GSSG, because an optimal condition for the PDI-catalyzed protein folding has relatively a reductive condition with a high proportion of GSH^21^. The results demonstrated that ERp46 introduces disulfide bonds more rapidly and promiscuously than PDI. However, considering that the redox balance in the ER^22^ is more oxidative than in these experimental conditions, we decided to use a buffer containing 2 mM GSH and 1 mM GSSG, which is more oxidative and mimics more closely physiological redox conditions.

High-performance liquid chromatography (HPLC) analysis showed that ERp46 in a buffer containing 2 mM GSH and 1mM GSSG converted fully the reduced form (R) of BPTI to several oxidized species within 1 min (Figures 1c and 1d). The HPLC elution profiles showed that a large number of small peaks, corresponding to off-pathway folding intermediates, were eluted around the R form during the initial stages of ERp46-catalyzed BPTI folding. Conversely, the HPLC elution profile of PDI-catalyzed BPTI folding revealed only a specific and well-defined elution peak near the R form, corresponding to an on-pathway disulfide-containing folding intermediate, in the early stage of folding.

After 5 min, the HPLC elution profiles showed more selective accumulation of on-path BPTI folding intermediates by PDI than by ERp46, and after 60 min, the HPLC elution profiles showed that PDI was more rapid than ERp46 in converting nearly all of the BPTI folding intermediates to the native folded state. These results are consistent with previous results obtained using a combination of ER oxidoreductin-1 α and Peroxiredoxin 4^15,19^, an upstream oxidase of the PDI family.

### Chaperone function of PDI/ERp46 against BPTI

The differences in the catalytic intermediates and speed of oxidative folding by PDI and ERp46 may be due to differences in their molecular recognition of BPTI along the folding pathway. To clarify these differences, we prepared the conformation of BPTI to a constitutive unfolded state by substituting its six Cys residues with Ser residues (BPTI all-Cys/Ser). In general, structural disulfide bonds stabilize the local and global conformations of folding intermediates and folded proteins by decreasing the entropy of the random coil unfolded state^23^. Figure 2a shows the positions of the six Cys/Ser substitutions in BPTI and the positions of the Cys residues in the crystal structure of native BPTI. The ^1^H-^15^N HSQC NMR spectrum of BPTI all-Cys/Ser showed narrow dispersion of the resonances, which is characteristic to unfolded proteins (Figure 2b).

**Figure 2.**
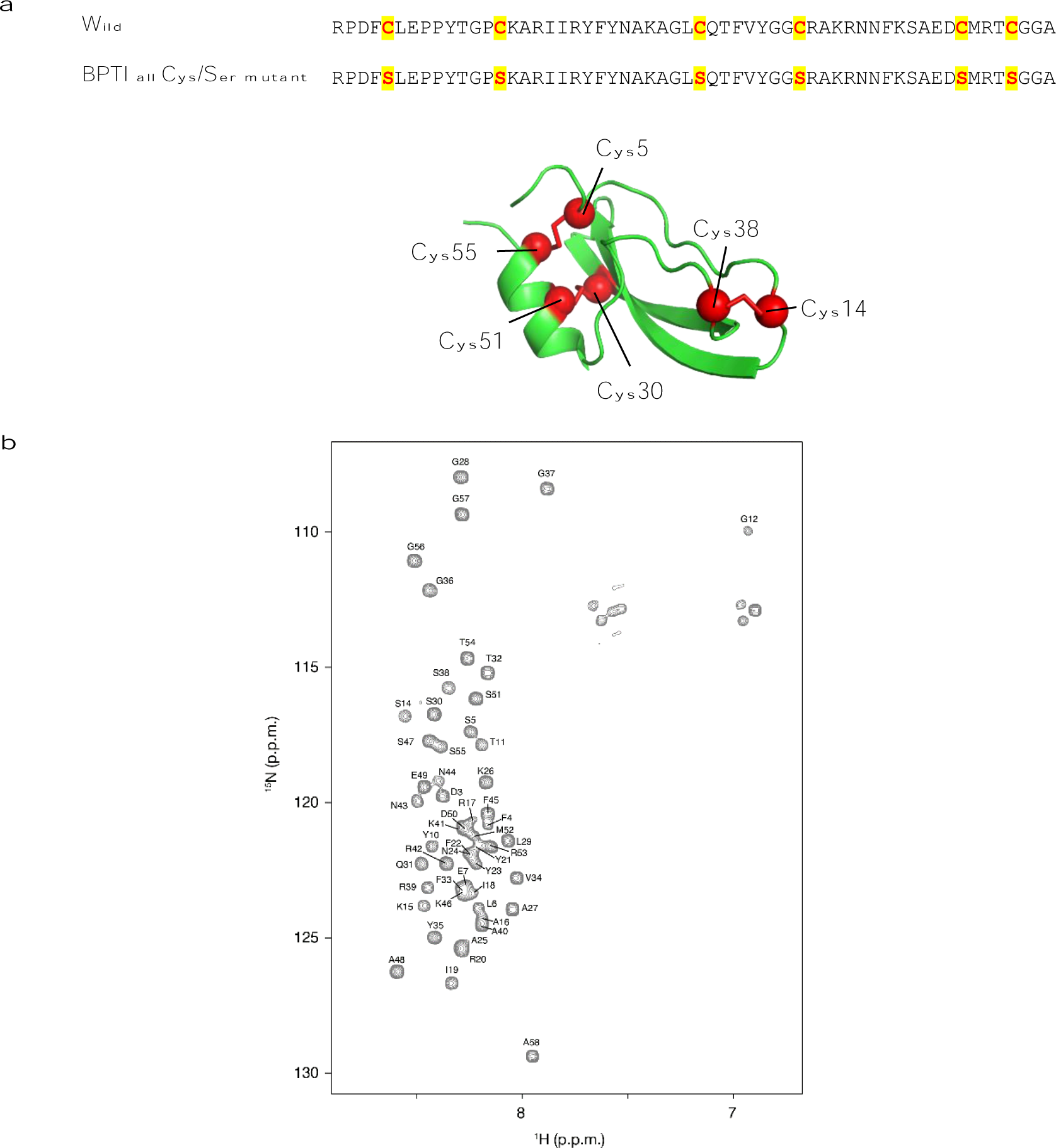
Cys/Ser substitutions in BPTI induce a constitutive unfolded state. (a) upper: Positions of the six Cys residues in the amino acid sequence of BPTI; lower: Positions of the Cys residues in the crystal structure of native BPTI. (b) 2D ^1^H-^15^N HSQC spectrum of BPTI with Cys/Ser substitutions

Turbidity measurements at 600 nm revealed that BPTI all-Cys/Ser aggregated when incubated at high concentration for 24 hours and co-incubation with the oxidized form of PDI but not the oxidized form of ERp46 considerably suppressed aggregation (Figure 3a). In fact, the oxidized form of ERp46 increased aggregation, perhaps suggesting that ERp46 forms co-aggregates with BPTI all-Cys/Ser. To confirm that oxidized PDI inhibited aggregation, we subjected the supernatants of the aggregation assays after centrifugation to non-reducing SDS-PAGE to separate the proteins by molecular weight. As shown in Fig. 3b, the chaperone function of PDI suppressed BPTI all-Cys/Ser aggregation, as evidenced by the increased intensity of the BPTI all-Cys/Ser monomer band migrating at the bottom of the gel compared to the lane without chaperone after 24 h incubation, whereas ERp46 actually promoted the aggregation of BPTI all-Cys/Ser, as evidenced by the continued decrease in the intensity of the monomeric BPTI all-Cys/Ser band. The bands migrating at approximately 55 kDa and 46 kDa correspond to oxidized PDI and oxidized ERp46, respectively. To verify whether PDI also suppresses BPTI all-Cys/Ser-ERp46 co-aggregation, we measured the turbidity of assays containing both PDI and ERp46. Although the theoretical values were calculated from the data in Figure 3a, the actual data unexpectedly showed that ERp46-mediated BPTI all-Cys/Ser aggregation was inhibited by PDI, suggesting that PDI suppresses ERp46-mediated aggregation of BPTI presumably at the initial stage of folding (Figure 3c). PDI and ERp46 together were previously shown to catalyze synergistically the oxidative folding of BPTI^19^. This may be interpreted as a new plausible scenario, in which PDI suppresses ERp46-client co-aggregation after ERp46 rapidly introduces disulfide bonds into the client.

**Figure 3.**
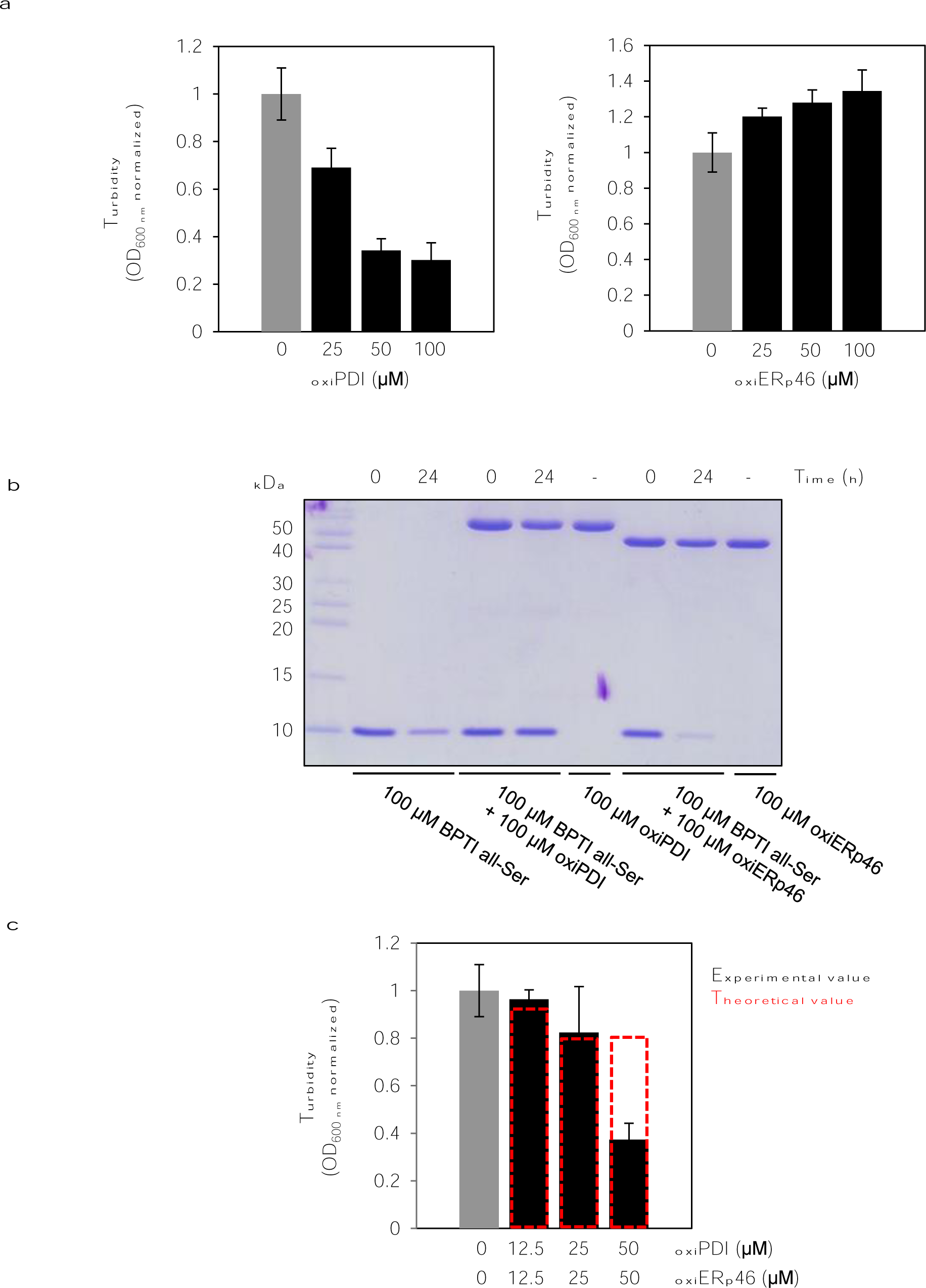
Chaperone function of PDI/ERp46 against Cys/Ser-mutated BPTI (BPTI all-ser) (a) Turbidity assays of 100 μM BPTI all-Cys/Ser with or without PDI/ERp46. Bars show mean ± SEM (n=3). (b) Non-reducing SDS-PAGE of BPTI all-ser after incubation with or without PDI/ERp46. (c) Turbidity assays of 100 μM BPTI all-ser after incubation with a mixture of PDI and ERp46. Bars show mean ± SEM (n=3).

### Client recognition differences between PDI and ERp46

To further understand the difference between the chaperone function and the catalytic function of disulfide bonds by PDI and ERp46 from the molecular recognition mechanism, the NMR spectrum of BPTI all-Cys/Ser was obtained in the absence and presence of ERp46 or PDI. Although ERp46 has high activity in introducing disulfide bonds into BPTI (Figure 1), the NMR signal of BPTI all-Cys/Ser remained almost unchanged by the addition of equimolar ERp46, suggesting that the interaction between BPTI all-Cys/Ser and ERp46 is weak (Figure 4). Conversely, the addition of PDI induced significant perturbations to the NMR signals of BPTI all-Cys/Ser, indicating relatively tight recognition of BPTI all-Cys/Ser by PDI (Figure 5).

**Figure 4.**
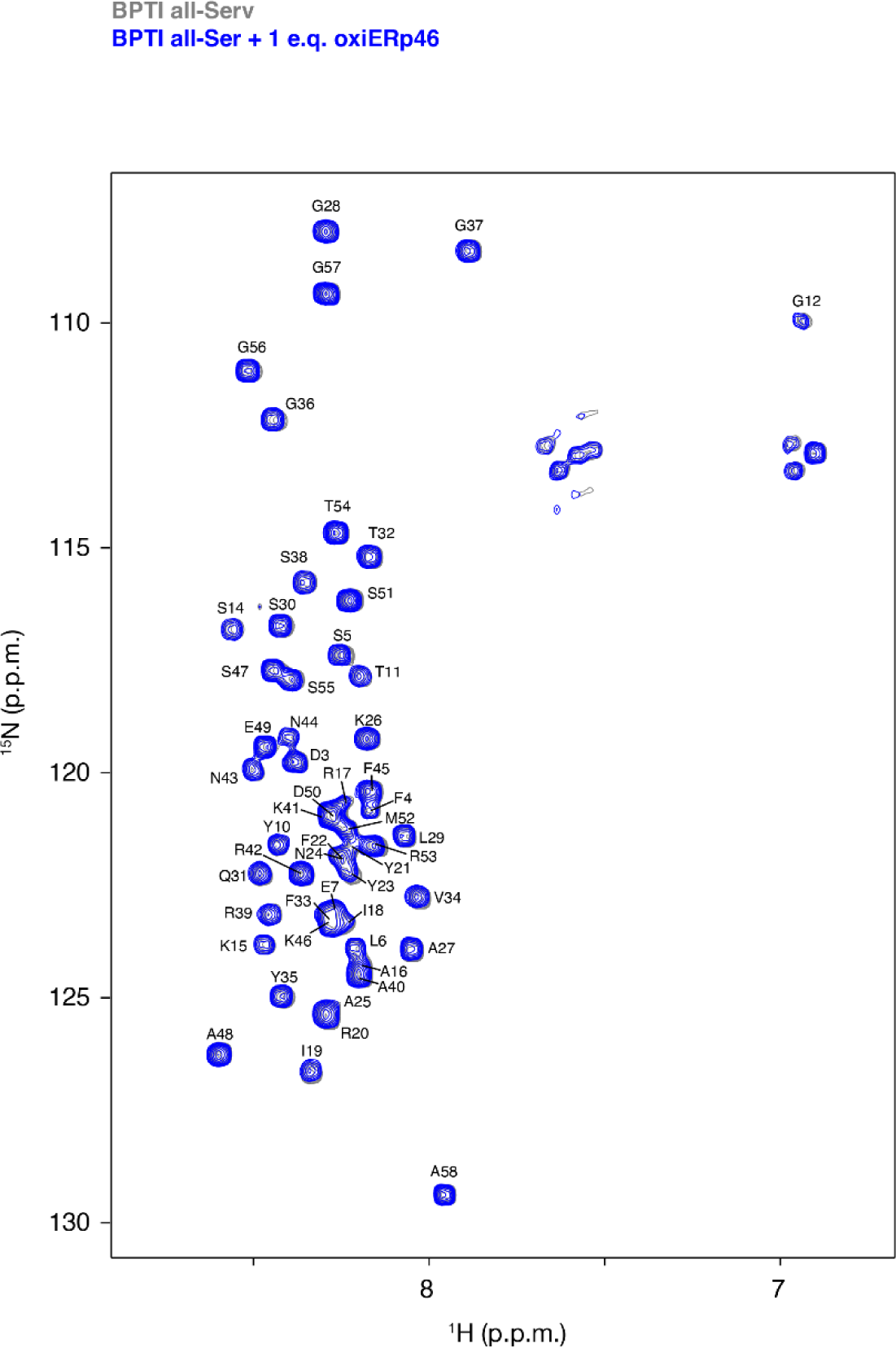
NMR spectra of BPTI all-ser after the addition of ERp46. ^1^H-^15^N HSQC spectra of BPTI all-ser in the absence (gray) and presence of an equimolar amount of ERp46 (blue).

**Figure 5.**
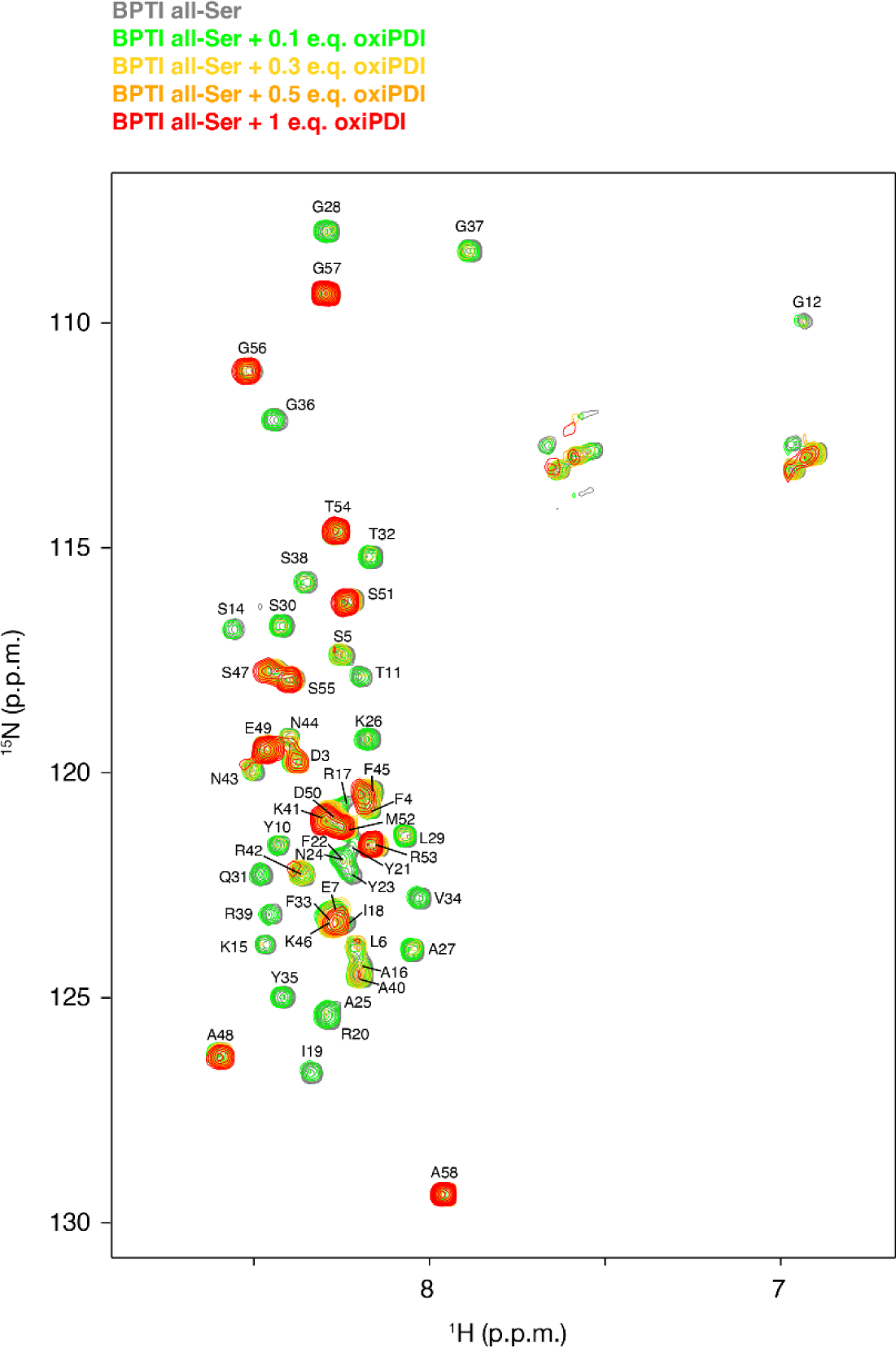
NMR spectra of BPTI all-ser after the addition of PDI. ^1^H-^15^N HSQC spectra of BPTI all-ser in the absence (gray) and presence of different amounts of PDI (0.1eq in green; 0.3 eq in yellow; 0.5eq in orange; 1.0 eq in red).

The residue-by-residue chemical shift and intensity changes of BPTI all-Cys/Ser induced by PDI showed that broad region of BPTI all-Cys/Ser encompassing from residues 7 to 40 are affected (Figure 6a). However, the chemical shift changes and intensity changes induced by ERp46 were much smaller than those by PDI (Figure 6b). These results highlighted the difference in the binding property of PDI and ERp46, where PDI shows relatively tight recognition of the client protein but ERp46 shows promiscuous interactions.

**Figure 6.**
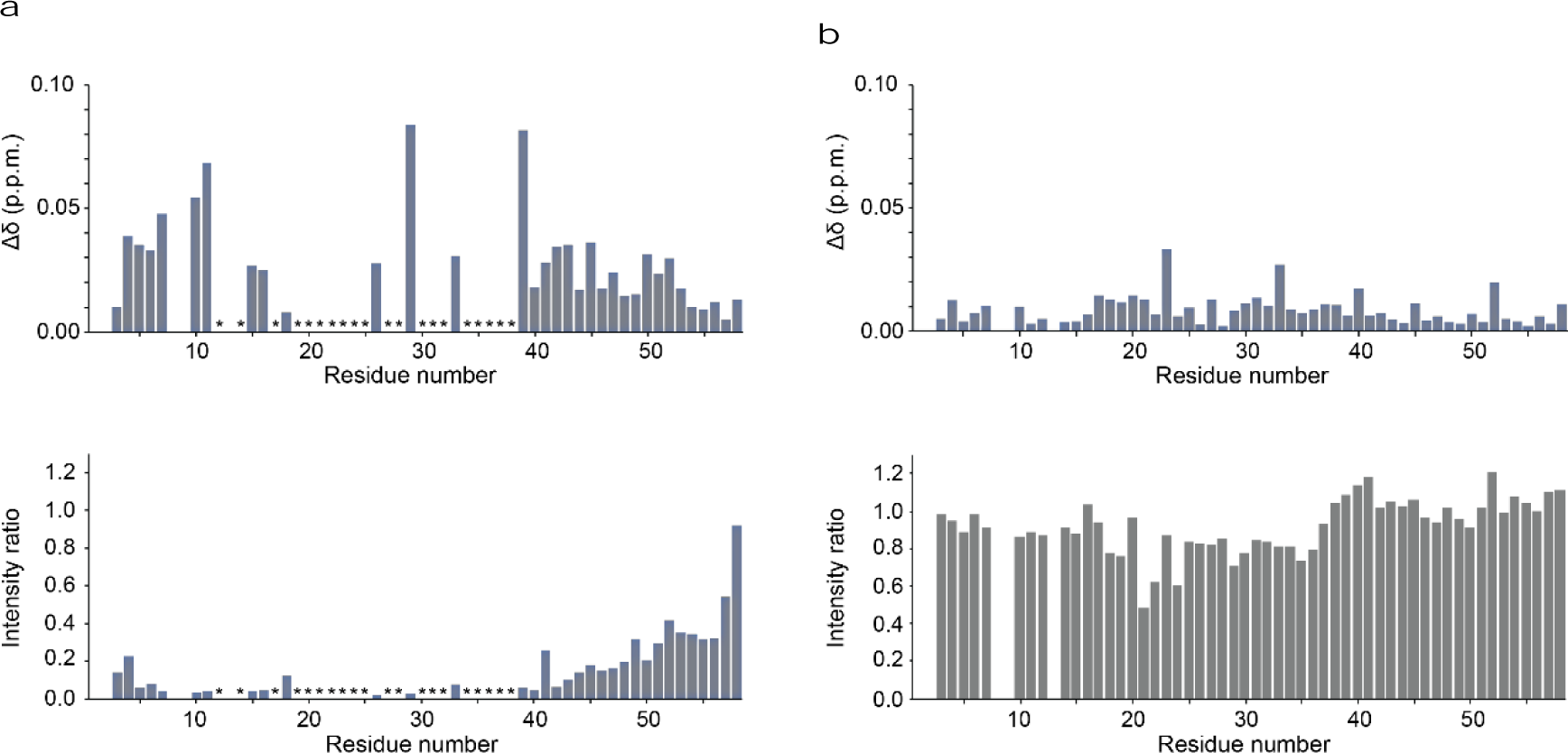
Client recognition differences between PDI and ERp46. (a) upper: Chemical shift changes of ^15^N-labeled Cys/Ser-mutated BPTI (BPTI all-ser) in the presence of PDI, lower: NMR spectrum intensity changes of ^15^N-labeled BPTI all-ser in the presence of PDI. The asterisks show indicates the residues whose signal disappeared by the addition of PDI. (b) upper: Chemical shift changes in ^15^N-labeled of BPTI all-ser in the presence of ERp46, lower: NMR spectrum intensity changes of ^15^N-labeled BPTI all-ser in the presence of ERp46.

In this study, we demonstrated that at the initial client folding stage, PDI binds specifically to a broad region from residues 7 to 40 within the unfolded BPTI within unfolded BPTI to function as a chaperone, while ERp46 interacts more broadly and weakly with unfolded BPTI, resulting in the rapid and promiscuous introduction of disulfide bonds. These different client recognition modes between PDI and ERp46 most likely play crucial roles in modulating enzymatic and chaperone activities. Client recognition differences between PDI family members likely contribute to determining whether disulfide bonds are introduced or aggregation of the unfolded client is suppressed during initial folding. Thus, the mammalian ER ensures concerted protein quality control due to the client recognition differences between PDI family members, resulting in the production of large quantities of multiple-disulfide bonded proteins.

## Acknowledgments

This research was funded by a JSPS KAKENHI grant (grant numbers 22H02205 (to MO), 22H02560 (to TS), 22K15278 (to MM), 22H04847 (to MM), Fund for the Promotion of Joint International Research (grant number 23KK0105 (to MO and MM), 20KK0156 (to TS)), a JSPS Grant-in-Aid for Transformative Research Areas (B) (grant number JP21H05095 (to MO) and 21H05094 (to TS)), the Japan Science and Technology Agency FOREST Program (grant number JPMJFR201F (to MO), JPMJFR204W (to TS)), HIRAKU-Global Program (MEXT’s “Strategic Professional Development Program for Young Researchers” (to MM)), the Takeda Science Foundation (to MO and TS), the Mochida Memorial Foundation for Medical and Pharmaceutical Research (to MO), the Naito Foundation (to MO and TS), the Uehara Memorial Foundation (to MO, SK, MM, and TS), the Terumo Life Science Foundation (to MO), Astellas Foundation for Research on Metabolic Disorders (to MO MM and TS), The Asahi Glass Foundation (to MO and TS), Mitsui Sumitomo Insurance Welfare Foundation (to MO), Daiichi Sankyo Foundation of Life Science (to MO), THE SUMITOMO FOUNDATION (to MO), and Cannon Foundation (to TS). This research was partially supported from Medical Research Center Initiative for High Depth Omics, and Joint Usage and Joint Research Programs, the Institute of Advanced Medical Sciences (IAMS), Tokushima University. Generous support was received from the FRIS CoRE, which is a shared research environment at Tohoku university. We thank Prof. Kenji Inaba for the kindly gift of PDI and ERp46 plasmids. Parts of the NMR experiments were performed at Hokkaido University Advanced NMR Facility, a member of NMR Platform.

## Author Contributions

TS and MO designed and supervised the current work. TS, KI, MM, HK, and SK performed NMR experiments. KI performed oxidative folding and chaperone assays under the supervision of SK and MO. MO wrote the first draft and TS revised the draft version. All the authors discussed the results, critically read the manuscript, and approved the final version for submission.

## Materials and Methods

### Materials

The recombinant aprotinin (bovine pancreatic trypsin inhibitor, #pro-285) was purchased from ProSpec-Tany TechnoGene Ltd, Israel.

### Expression and purification of recombinant proteins

cDNA encoding BPTI was subcloned into the NdeI and EcoRI sites of the pET17b vector (Novagen, Germany). The Cys/Ser-substituted BPTI mutant (BPTI all-Cys/Ser) was constructed using the PrimeSTAR Mutagenesis Basal Kit (Takara Bio, Japan). BPTI all-Cys/Ser was overexpressed in *Escherichia coli* strain BL21(DE3) by culturing at 37°C overnight. The cells were disrupted in buffer A (50 mM Tris-HCl, pH 8.1, and 300 mM NaCl) containing 1 mM phenylmethylsulfonyl fluoride using a homogenizer (Sonics and Materials, USA). After centrifugation of the homogenized lysate, recombinant BPTI all-Ser protein was obtained in the form of inclusion bodies. The inclusion bodies were dissolved in 100 mM Tris-HCl (pH 8.0) containing 8 M urea and 20 mM dithiothreitol at 50°C for 1 h. After centrifugation, the supernatant was loaded onto a COSMOSIL 5C18 column (Nacalai Tesque, Japan), and BPTI all-Ser was eluted with 80% CH_3_CN in 0.05% TFA. The eluted samples were further purified by RP-HPLC (GL Science, Japan) equipped with a COSMOSIL 5C18-AR-II column (Nacalai Tesque, Japan) using a linear gradient of CH_3_CN in 0.05% TFA. Purified samples were lyophilized.

Plasmids expressing human PDI and ERp46 without their signal sequences were constructed previously^16,19,24^. Human PDI and ERp46 were overexpressed in *Escherichia coli* strain BL21(DE3) containing each plasmid by the addition of 0.5 mM isopropyl-β-D-thiogalactopyranoside and culturing at 20°C overnight. After centrifugation, the cells were disrupted in buffer A (50 mM Tris-HCl, pH 8.1, and 300 mM NaCl) using an ultrasonic homogenizer (Branson Sonifier SFX250, Emerson, USA). After centrifugation, the supernatant was loaded onto a Ni-NTA agarose column (Wako, Japan). The column was washed with buffer A containing 20 mM imidazole, and the recombinant proteins were eluted with buffer A containing 200 mM imidazole. The eluted proteins were applied to a Resource Q anion exchange column (Cytiva, USA) pre-equilibrated with 50 mM Tris-HCl (pH 8.1) and eluted with a linear gradient of 0–500 mM NaCl. The proteins were further purified by size-exclusion chromatography on a Superdex 200 Increase column (Cytiva, USA) pre-equilibrated with buffer A. Oxidized PDI and ERp46 were prepared by incubating purified PDI and ERp46 with 10 mM diamide as an oxidant on ice for 60 min. After incubation, the oxidant was removed by a Superdex 200 Increase column (Cytiva, USA) chromatography. Protein concentrations were determined using the Protein Assay BCA Kit (Nacarai Tesque, Japan).

### Oxidative folding assay

Reduced and denatured BPTI (30 μM) was incubated with/without 1 μM PDI or ERp46 in 100 mM Tris-HCl (pH 7.5) containing 300 mM NaCl, 2 mM GSH, and 1 mM GSSG at 30°C, as described previously^15,17,19,25^. The reaction was quenched with an equivalent volume of 1 M HCl at the indicated time points and separated by RP-HPLC on a TSKgel Protein C4-300 column (Tosoh Bioscience, Japan) with monitoring at 229 nm. The identities of the resulting peaks were confirmed by MALDI-TOF/MS analysis as described previously^25,26^.

### Chaperone assay

BPTI all-Ser (100 μM) was incubated with/without various concentrations (0–100 μM) of oxidized PDI and/or ERp46 in 50 mM HEPES-NaOH (pH 7.0) and 500 mM NaCl at 37°C for 24 h. BPTI aggregation was monitored as turbidity at 600 nm using a UH3900 spectrophotometer^11,12^ (Hitachi-Hightech, Japan). For SDS-PAGE analysis, all samples were centrifuged and the supernatants were separated by non-reducing SDS-PAGE. The gel bands were visualized by the 2,2,2-trichloroethanol in-gel method^27^ using a ChemiDoc imager (Bio-Rad, CA).

### NMR experiment

Isotopically-labeled BPTI all-Ser was prepared at a concentration of 0.1 mM in 50 mM HEPES-NaOH (pH 7.0), 200 mM NaCl and 10% ^2^H_2_O. For resonance assignment, ^13^C/^15^N-labeled BPTI all-Ser was subjected to NMR measurements on a Bruker Avance Neo 800 MHz NMR spectrometer equipped with a TCI cryogenic probe. Backbone resonance assignment for BPTI all-Ser was carried out using the following set of spectra; ^1^H–^15^N selective optimized flip angle short transient (SOFAST)-heteronuclear multiple quantum correlation (HMQC)^28^, constant time ^1^H-^13^C heteronuclear single quantum correlation (HSQC), HNCO, HNCA, HN(CO)CA, HNCACB, CBCA(CO)NH, and C(CO)NH spectra at a sample temperature of 10°C. The spectra were processed using Topspin. Resonance assignment was performed with the SPARKY program (T. D. Goddard and D. G. Kneller, SPARKY 3, University of California, San Francisco; https://www.cgl.ucsf.edu/home/sparky/), which resulted in the assignment of 98% of the backbone amide resonances. For interaction analysis with PDI family proteins, ^15^N-labeled BPTI all-Ser was subjected to NMR measurements on a Bruker Avance III 500 MHz NMR spectrometer equipped with a BBO probe at a sample temperature of 10°C. ^1^H–^15^N SOFAST-HMQC spectra of ^15^N-labeled BPTI all-Ser were acquired in the absence and presence of oxidized PDI at 10, 30, 50, and 100 µM or ERp46 at 100 µM. The spectra were processed using the NMRPipe program^29^ and analyzed using the Olivia program (https://github.com/yokochi47/Olivia).

